# Genome-scale analysis of cellular restriction factors that inhibit transgene expression from adeno-associated virus vectors

**DOI:** 10.1101/2022.07.13.499963

**Authors:** Ashley M Ngo, Andreas S Puschnik

## Abstract

Adeno-associated virus (AAV) vectors are one of the leading platforms for gene delivery for the treatment of human genetic diseases, but the antiviral cellular mechanisms that interfere with optimal transgene expression are incompletely understood. Here, we performed two genome-scale CRISPR screens to identify cellular factors that restrict transgene expression from recombinant AAV vectors. Our screens revealed several components linked to DNA damage response, chromatin remodeling and transcriptional regulation. Inactivation of the Fanconi Anemia gene FANCA, the Human Silencing Hub (HUSH) associated methyltransferase SETDB1 and the gyrase, Hsp90, histidine kinase and MutL (GHKL)-type ATPase MORC3 led to increased transgene expression. Moreover, SETDB1 and MORC3 knockout improved transgene levels of several AAV serotypes as well as other viral vectors, such as lentivirus and adenovirus. Finally, we demonstrated that inhibition of FANCA, SETDB1 or MORC3 also enhanced transgene expression in human primary cells, suggesting that these could be physiologically relevant pathways that restrict AAV transgene levels in therapeutic settings.

**IMPORTANCE:** Recombinant AAV (rAAV) vectors have been successfully developed for the treatment of genetic diseases. The therapeutic strategy often involves the replacement of a defective gene by expression of a functional copy from the rAAV vector genome. However, cells possess antiviral mechanisms that recognize and silence foreign DNA elements thereby limiting transgene expression and its therapeutic effect. Here, we utilize a functional genomics approach to uncover a comprehensive set of cellular restriction factors that inhibit rAAV-based transgene expression. Genetic inactivation of selected restriction factors increased rAAV transgene expression. Hence, modulation of identified restriction factors has the potential to enhance AAV gene replacement therapies.

## INTRODUCTION

Adeno-associated virus (AAV) provides a powerful and versatile platform for gene delivery to treat a variety of human genetic diseases. AAV is a small single-stranded DNA virus with a genome size of ~4.7 kb from the *Parvoviridae* family. Its genome consists of two open reading frames, *rep* and *cap*, which are flanked by inverted terminal repeat (ITR) sequences (1, 2). The ITRs are the only cis-acting elements required for genome packaging, which enables the generation of recombinant AAV (rAAV) vectors containing a reporter or therapeutic transgene in place of the endogenous *rep* and *cap* genes (3). rAAV vectors have been clinically approved for the treatment of monogenic diseases, such as lipoprotein lipase deficiency, Leber congenital amaurosis and spinal muscular atrophy, and there are over 150 ongoing clinical studies for additional rAAV-based therapies (4).

Despite the recent clinical successes, rAAV-based gene therapies face several challenges: (i) Large-scale manufacturing of rAAV for high dose administration (>10^14^ viral particles per patient), is expensive and difficult; (ii) immunological barriers such as the presence of pre-existing neutralizing antibodies against AAV or the triggering of a robust humoral immune response upon treatment prevent (repeated) administration; (iii) hepatic and neuronal genotoxicity has been observed in a number of high-dose administrations; (iv) transgene expression can be lost over time due to the clearance of transduced cells through a cytotoxic T cell response and/or potential silencing of transgene expression (4–6). Increased efficiency in rAAV transgene expression could therefore allow for a reduction in the therapeutic doses needed, which consequently could decrease costs, elicit fewer neutralizing antibodies, reduce the risk of genotoxicity, and lower AAV capsid-directed T lymphocyte-mediated cytotoxicity. However, the precise host cellular pathways that restrict rAAV transgene expression are incompletely understood.

## RESULTS

### Genome-wide CRISPR screens identify rAAV restriction factors

To identify cellular restriction factors of rAAV transgene expression that are likely to be conserved across multiple cell types, we performed two parallel genome-wide screens using CRISPR knockout (KO) in A549 human lung adenocarcinoma cells and CRISPR interference (CRISPRi) in K562 myelogenous leukemia cells. We used a non-saturating multiplicity of infection (moi) so that inhibition of a repressor would lead to an increase of expression of a fluorescent reporter transgene. Based on rAAV serotype 2 (rAAV2) moi titrations, we used 10,000 viral genomes (vg)/cell for the A549 screen and 50,000 vg/cell for the K562 screen (Figure S1A). We performed two rounds of rAAV transduction and FACS-based enrichment of the top 10-20% fluorescent CRISPR library cells followed by analysis of the guide RNA (gRNA) representation via next-generation sequencing (Figure 1A). Analysis with MaGeCK, a computational tool that uses Robust Rank Aggregation (RRA) to identify positively or negatively selected genes, resulted in significant enrichment of 20 genes in the A549 screen and of 88 genes in the K562 screens with a cutoff of -log(enrichment score) >5 (Figure S1B and S1C, Table S1 and S2). Nearly all individual gRNAs were enriched for the top 15 genes of both screens, suggesting that the increased transgene expression phenotypes resulting from the genetic perturbations were robust (Figure S1D). Comparison of the enrichment scores from the A549 and K562 CRISPR screens highlighted that several Fanconi Anemia (FA) genes, the histone methyltransferase SETDB1, the GHKL-type ATPase MORC3, the SUMO Interacting Motifs Containing 1 (SIMC1), the DNA damage response gene ATM and additional genes (SMCHD1, MTF1, DCLRE1A, DAXX, FAM208A, ZRANB2) were enriched in both datasets (Figure 1B). STRING analysis on the top 100 hits from both screens, identified functional gene clusters and cellular pathways, including the FA pathway, the Human Silencing Hub (HUSH) complex containing the SETDB1 histone methyltransferase, the SMC5-SMC6 complex, the CCR4-NOT complex, and genes linked to DNA damage repair and chromatin regulation (Figure 1C, Table S3).

**Figure 1.**
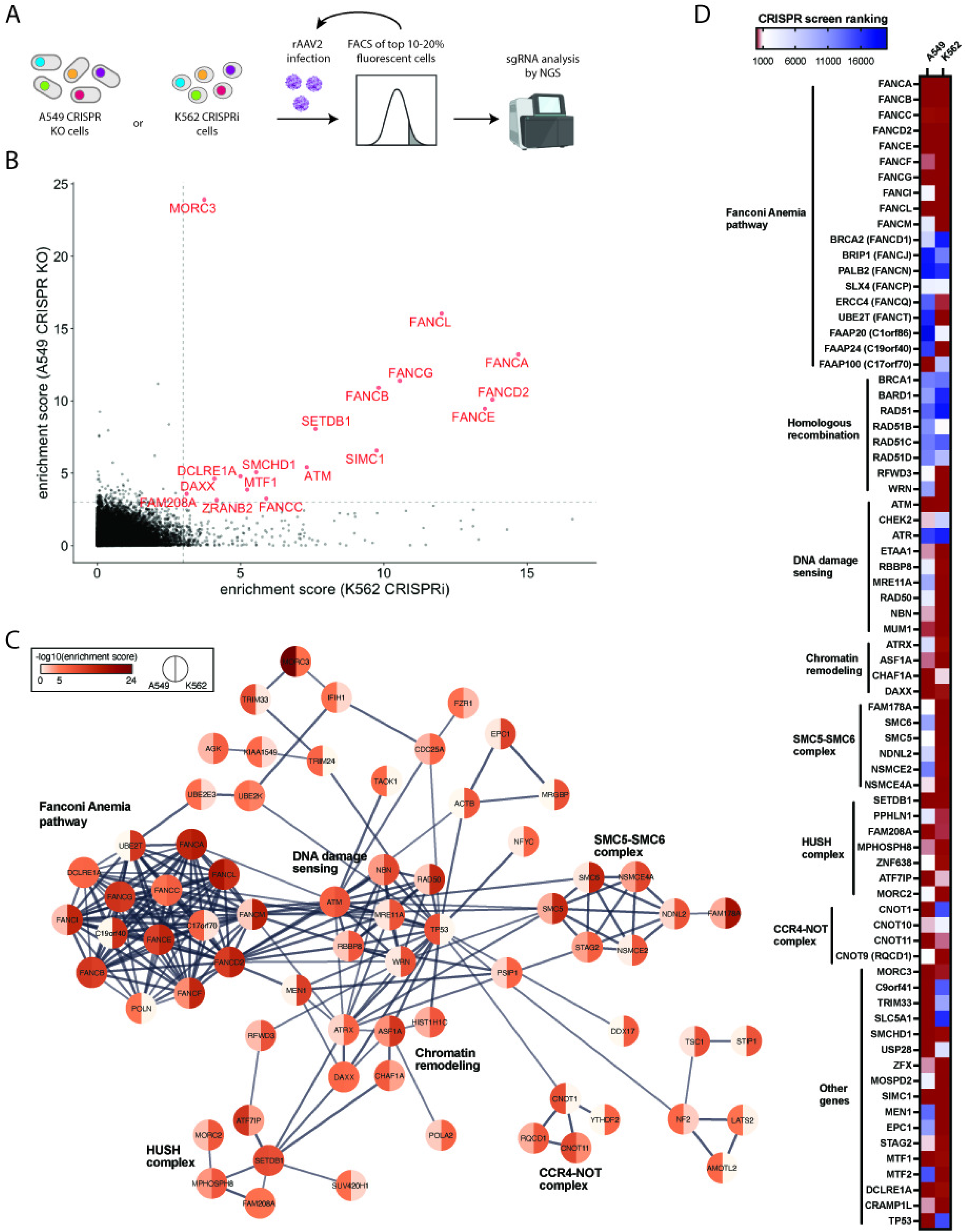
CRISPR screens reveal cellular factors that restrict rAAV transgene expression. (A) Schematic of genome-wide CRISPR screens using CRISPR KO in A549 human lung adenocarcinoma cells and CRISPRi in K562 myelogenous leukemia cells. Cells were transduced with rAAV2 expressing a fluorescent reporter gene (GFP or tdTomato) and the top 10-20% fluorescent cells were sorted. After 5-7 days, a second round of transduction and sorting was performed before cells were collected for analysis of sgRNA enrichment by next-generation sequencing. (B) MaGeCK gene enrichment scores of the A549 CRISPR KO screen (y-axis) vs. MaGeCK gene enrichment scores of K562 CRISPRi screen (x-axis). Genes with -log10 (enrichment score) > 3 in both screens are highlighted. (C) STRING network for the top 100 hits from both the A549 and the K562 CRISPR screens. Genes were clustered based on high-confidence interactions (STRING confidence cutoff 0.7) and the major gene network is displayed. CRISPR screen enrichment scores for each gene are shown using a color scale. Line thickness indicates confidence of interaction. (D) Heatmap of gene rankings in A549 and K562 CRISPR screens for selected genes related to DNA damage response and other cellular processes.

The FA pathway is a biochemical network of 19 genes which are responsible for DNA repair of interstrand crosslinks and stabilization of stalled replication forks, which can impede DNA replication and transcription (7). Most FA core complex components (FANCA-FANCM), which are responsible for lesion recognition and binding, were robustly enriched in both rAAV screens, while the FA factors mediating lesion resolution downstream and the homologous recombination machinery were not or only partially enriched (Figure 1D). Another DNA damage repair factor that scored highly in both screens is the serine/threonine protein kinase ATM (Figure 1B and 1D). ATM activates checkpoint signaling upon double strand breaks, apoptosis and genotoxic stresses, thereby acting as a DNA damage sensor (8). Loss of the FA pathway and ATM is synthetic lethal, suggesting closely linked cellular roles (9). Besides DNA damage response genes, many factors were linked to chromatin and transcriptional regulation (Figure 1C and 1D). The histone methyltransferase SETDB1 is recruited by the HUSH complex, composed of FAM208A (also known as TASOR), MPHOSPH8 and PPHLN1, to heterochromatic loci and foreign DNA elements, such as retroviruses and transposons, to mediate transcriptional repression (10, 11). Other HUSH associated factors (MORC2, ZNF638 and ATF7IP) required for recruitment and full epigenetic silencing activity were also enriched in both screens (12–14). The SMC5-SMC6 complex, highly ranked in the K562 screen, was previously shown to play a role in silencing of extrachromosomal HBV DNA as well as unintegrated HIV-1 DNA (Figure 1C and 1D) (15, 16). Finally, MORC3 is a nuclear matrix protein with ATPase activity and has been linked to gene silencing, including of viral pathogens (17–19). Overall, the screens identified numerous candidate restriction factors for rAAV transgene repression, which have canonical roles in DNA damage repair and chromatin regulation.

### The Fanconi Anemia pathway suppresses transgene expression from rAAV vectors

The FA pathway has to our knowledge not been functionally associated with the regulation of AAV transgene expression. To validate the impact of the FA pathway on rAAV transgene expression, we transduced three sets of FANCA expressing and matched FANCA deficient cells. Dual gRNA/Cas9-mediated deletion of the N-terminus of FANCA in UM-SCC-01 cells significantly increased the percentage of rAAV2-tdTomato positive cells (rAAV2+) as well as their median fluorescence intensity (MFI) relative to WT cells (Figure 2A and 2B). This effect was reversed in FANCA transgene-complemented derivatives of the KO cells (Figure 2A and 2B). Furthermore, transduction of mouse ear fibroblasts (MEFs) derived from WT or FANCA KO 129S4/SvJaeSor mice showed a similar increase of rAAV transgene expression in the FANCA-deficient background (Figure 2C and 2D). Lastly, we obtained FA patient derived head and neck squamous cell carcinoma cells (“974 cells”), which harbor a FANCA frameshift mutation, as well as corresponding FA patient cells that were lentivirally complemented with an intact FANCA cDNA copy (20). Complementation with FANCA cDNA led to a substantial decrease of rAAV transgene expression relative to the unaltered FA patient derived 974 cells (Figure 2E and 2F). As the FA pathway plays a role in DNA repair, we next tested whether mutations in FANCA affect the stability of rAAV genomes in transduced cells. Quantitative PCR (qPCR) showed that intracellular rAAV genome copy numbers were not significantly different in FANCA KO relative to WT or FANCA trans-complemented UM-SCC1 cells (Figure 2G). Together, these data support that the FA pathway plays a role in the transgene expression from rAAV vectors, independent of rAAV genome stability.

**Figure 2.**
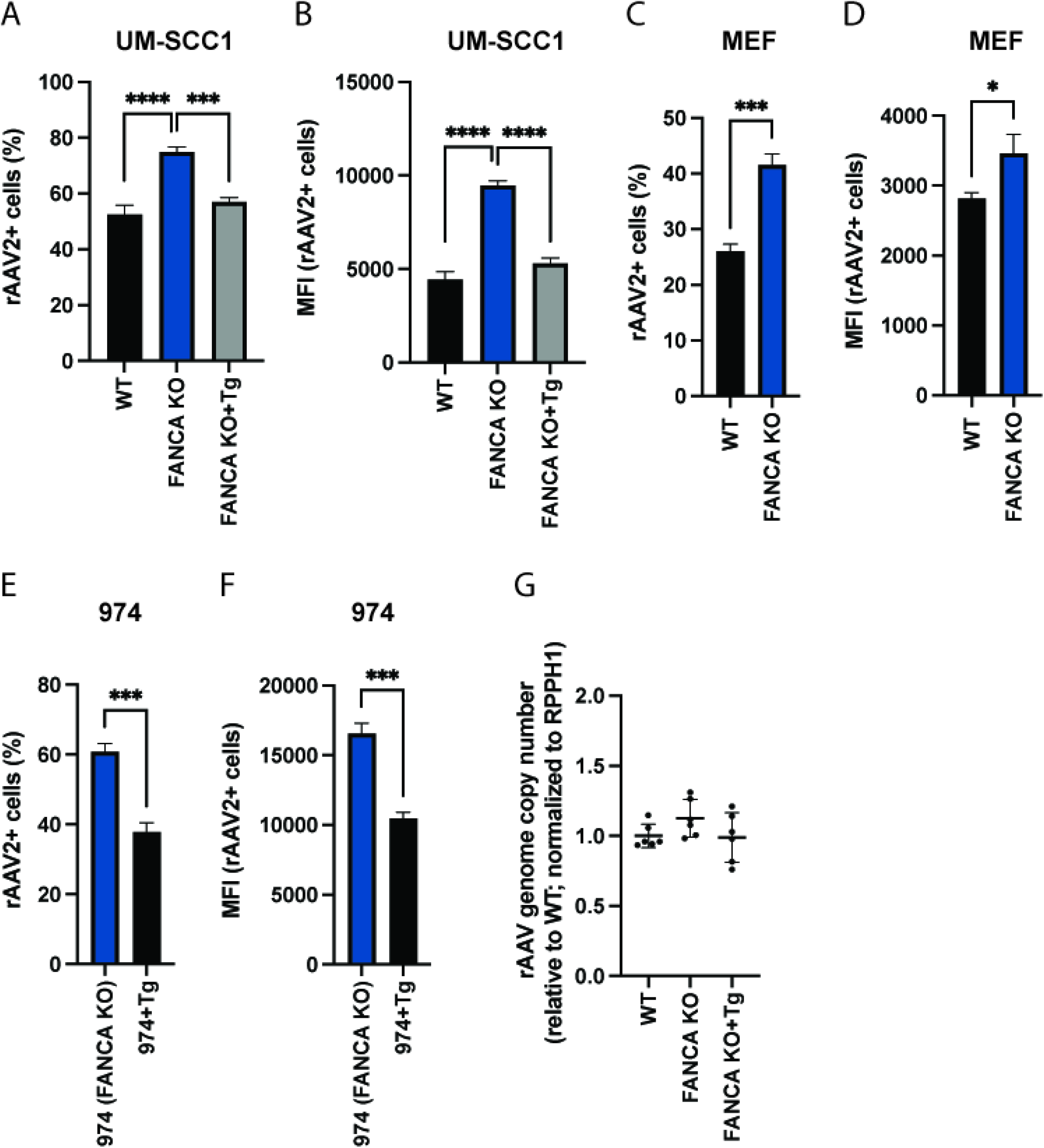
The Fanconi Anemia pathway represses rAAV transgene expression. (A) rAAV2-tdTomato infection rate or (B) MFI of human WT, FANCA KO and KO UM-SCC1 cells complemented with FANCA transgene (Tg). Cells were infected with rAAV2 (moi=10,000) for 48h and tdTomato expression was analyzed by flow cytometry. (C) rAAV2-tdTomato infection rate or (D) MFI of mouse ear fibroblasts (MEF) derived from WT or FANCA KO 129S4/SvJaeSor mice. Cells were infected with rAAV2 (moi=100,000) for 24h and tdTomato expression was analyzed by flow cytometry. (E) rAAV2-tdTomato infection rate or (F) MFI of FA patient-derived head and neck squamous cell carcinoma cells (974 cells), which harbor a FANCA frameshift mutation, as well as of 974 cells that were complemented with FANCA transgene (Tg). Cells were infected with rAAV2 (moi=10,000) for 48h and tdTomato expression was analyzed by flow cytometry. (G) Quantification of intracellular rAAV genome copies. Transduced cells (moi=10,000) were harvested after 48h, DNA was extracted and AAV and cellular RPPH1 DNA copy numbers were determined by qPCR. Data is combined from 2 independent experiments with 3 biological replicates each. All FACS data are shown as the mean ± s.d. from biological triplicates. P-values were determined by ANOVA with post-hoc Tukey’s test for (A)-(B) and unpaired t-test for (C)-(F), and are defined as follows: ns: non-significant; *: <0.05; **: <0.01; ***: <0.001; ****: <0.0001.

### Knockout of MORC3 and SETDB1 enhances rAAV transgene expression

Next, we sought to validate the impact of the (GHKL)-type ATPase MORC3 and the histone methyltransferase SETDB1 on rAAV as previous reports have shown antiviral functions for other DNA viruses (10, 14, 18, 19). We generated two clonal A549 KO cells per gene. The MORC3 KO clones harbored frameshift mutations in exon 1, while the SETDB1 KO clones contained a frameshift or large deletion (−21bp) in exon 2 (Figure S2A). The generated KO cell lines did not show any growth defects (Figure S2B). Infection of KO cells with rAAV2-tdTomato showed a substantial increase in rAAV2+ cells (by 16-19% in MORC3 KO and by 22-23% in SETDB1 KO cells 3 days post-infection (dpi)) as well as in MFI (by 2.7-3.3x in MORC3 KO and by 5.4-6.8x in SETDB1 KO cells 3dpi) relative to WT cells (Figure 3A, 3B and S3). rAAV transgene expression remained higher in KO vs WT cells over 8 days. Signal decreased in all conditions at later timepoints likely due to the dilution of episomal rAAV DNA during cell proliferation. Lentiviral complementation of MORC3 KO cells with MORC3 cDNA lowered the percentage of rAAV+ cells and MFI to levels observed in WT cells (Figure 3C and 3D), supporting that rAAV transgene expression is indeed modulated by MORC3 activity. Similarly, complementation of SETDB1 KO cells with SETDB1 cDNA decreased both the percentage of tdTomato+ cells (albeit to a lesser degree) and MFI (Figure 3E and 3F). Next, we tested whether deletion of MORC3 or SETDB1 allowed for achieving comparable rAAV transduction rates with reduced rAAV moi. Across a range of 100 to 100,000 vg/cell, MORC3 and SETDB1 KO cells exhibited significantly higher transgene expression levels than WT cells, demonstrating that only ~one-tenth of the rAAV amount was needed in KO cells to achieve similar transgene expression levels as in WT cells (Figure 3G and 3H).

**Figure 3.**
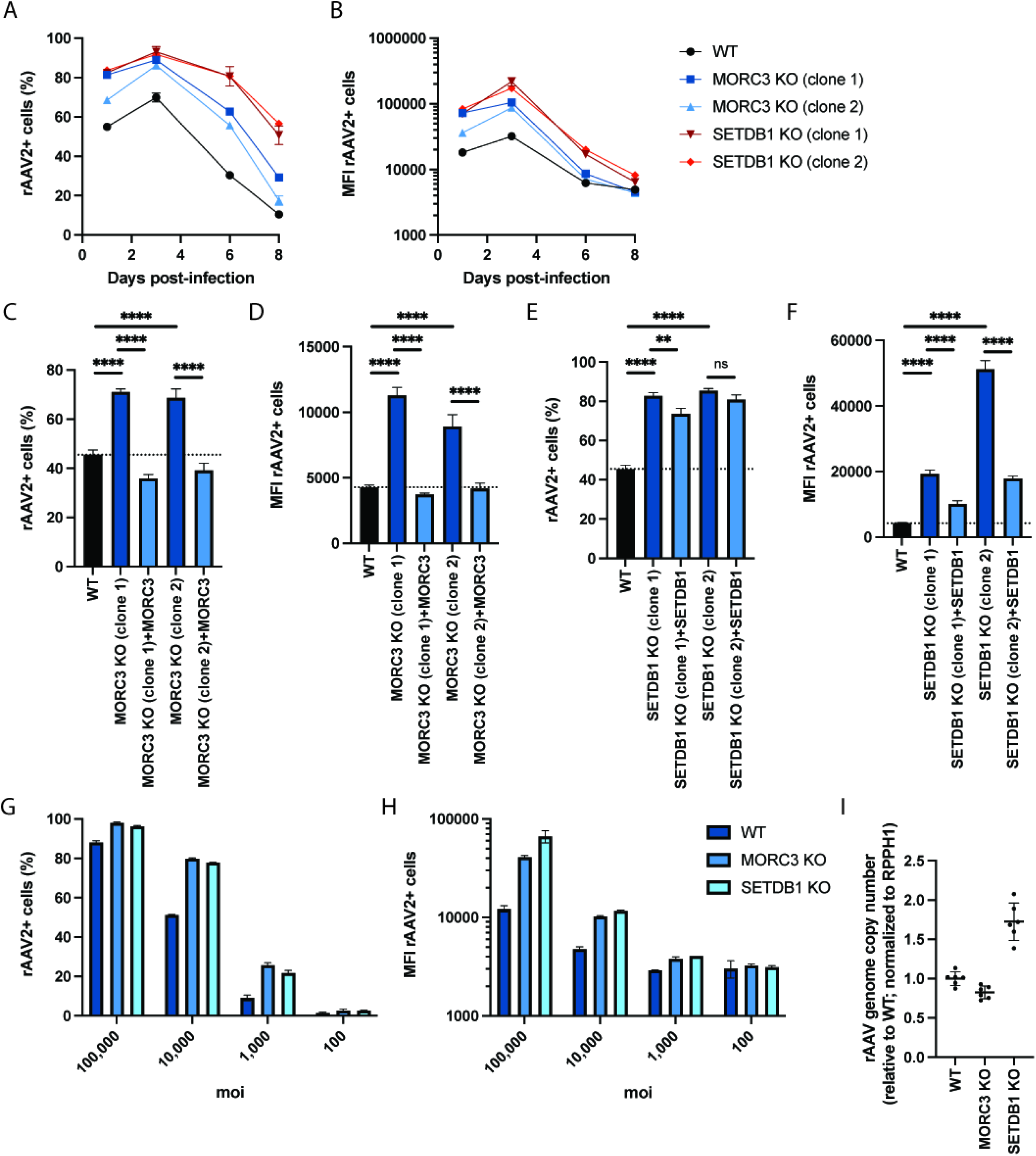
Knockout of MORC3 and SETDB1 increases rAAV2 transgene expression. (A) Percentage or (B) MFI of rAAV2-positive cells over time. WT, MORC3 KO or SETDB1 KO A549 cells were infected with tdTomato-expressing rAAV2 (moi=10,000) and cells were measured by flow cytometry at day 1, 3, 6 and 8 post-infection. (C) Percentage or (D) MFI of rAAV2-positive WT, MORC3 KO or cDNA-complemented MORC3 KO cells. Cells were infected with tdTomato-expressing rAAV2 (moi=10,000) and cells were measured by flow cytometry 24 hpi. (E) Percentage or (F) MFI of rAAV2-positive WT, SETDB1 KO or cDNA-complemented SETDB1 KO cells. Cells were infected with tdTomato-expressing rAAV2 (moi=10,000) and cells were measured by flow cytometry 24 hpi. (G) Percentage or (H) MFI of rAAV2-positive WT, MORC3 KO or SETDB1 KO cells infected with different multiplicities of infection (moi) and measured by flow cytometry 24 hpi. (I) Quantification of intracellular rAAV genome copies in WT A549, MORC3 KO (clone 1) and SETDB1 KO (clone 1). Transduced cells (moi=10,000) were harvested after 48h, DNA was extracted and AAV and cellular RPPH1 DNA copy numbers were determined by qPCR. Data is combined from 2 independent experiments with 3 biological replicates each. FACS data are shown as the mean ± s.d. from biological triplicates (A-F) or from biological duplicates (G-H). P-values were determined by ANOVA with post-hoc Tukey’s test and are defined as follows: ns: non-significant; *: <0.05; **: <0.01; ***: <0.001; ****: <0.0001.

To explore whether the restriction factors act on rAAV genome stability or transcriptional regulation, we measured whether rAAV genome copy numbers differed between WT, MORC3 KO and SETDB1 KO cells using qPCR. While the rAAV2 genome copy number was slightly lower in MORC3 KO cells 48h post-transduction, it was increased by 1.7x in SETDB1 KO cells (Figure 3I). Additional quantification of transgene mRNA levels via RT-qPCR showed that expression was moderately increased by 1.1-1.3x in MORC3 KO cells and by >2x in SETDB1 KO cells (Figure S4A). Together, these data indicate that MORC3 likely modulates rAAV transgene expression independently of affecting viral vector genome copies as MORC3 KO exhibited higher tdTomato levels despite lower rAAV genome copies. By contrast, deletion of the histone methyltransferase SETDB1 increased rAAV genome copies, which could largely explain the increase in tdTomato fluorescent levels. However, SETDB1 KO led to an even larger increase in transgene mRNA levels, which means that there could be additional restriction on transcriptional level.

Different rAAV serotypes are used in research and clinical applications, often depending on the target cell type or tissue. We therefore assessed whether MORC3 and SETDB1 also affected transgene expression from different rAAV serotypes. We infected cells with rAAV1, rAAV5 and rAAV6 vectors, which only lowly expressed GFP transgene in WT A549 cells. By contrast, GFP expression was drastically increased in MORC3 and SETDB1 KO cells (Figure 4A and 4B). Moreover, rAAV vector genomes can be single-stranded (ssAAV), which require second-strand synthesis prior to gene expression (as were all rAAV vectors used above), or they can be constructed with self-complementary genomes (scAAV). scAAV vectors form intra-molecular dsDNA and thus do not require cell-mediated synthesis of the second strand. We observed that presence of MORC3 and SETDB1 also restricted transgene expression from scAAV genomes, similarly as for ssAAV vectors, which suggests that these cellular restriction factors do not affect processes in the second-strand synthesis (Figure 4C and 4D).

**Figure 4.**
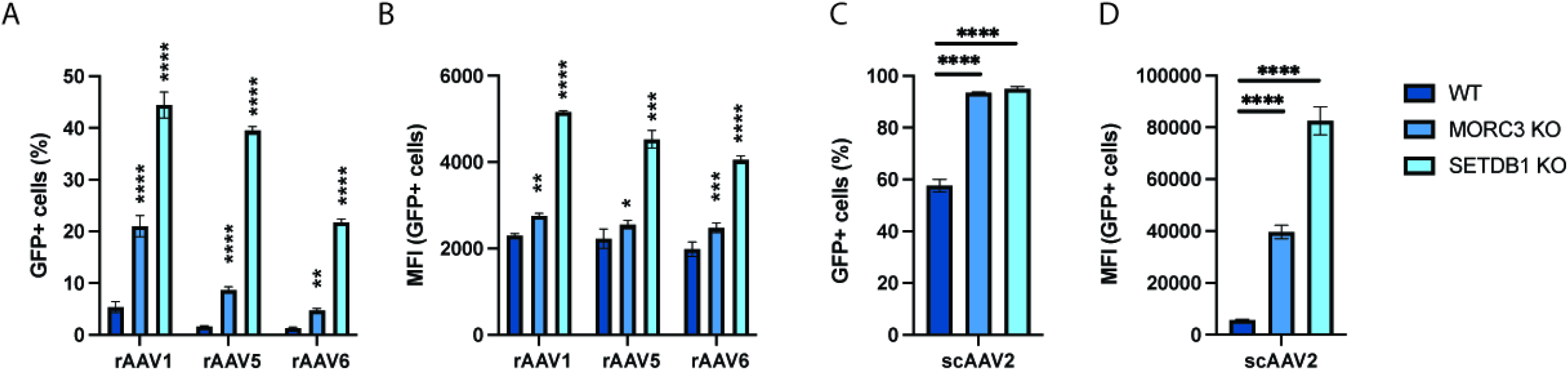
MORC3 and SETDB1 restrict transgene expression from different rAAV serotypes as well as self-complementary rAAV vectors. (A) Percentage or (B) MFI of WT, MORC3 KO or SETDB1 KO A549 cells infected with rAAV serotypes 1, 5 or 6 (moi=50,000) and analyzed by flow cytometry at 3 dpi. (C) Percentage or (D) MFI of WT, MORC3 KO or SETDB1 KO A549 cells infected with scAAV serotype 2 (moi=10,000) and analyzed by flow cytometry at 3 dpi. All data are shown as the mean ± s.d. from biological triplicates. P-values were determined by ANOVA with post-hoc Tukey’s test for each rAAV and are defined as follows: ns: non-significant; *: <0.05; **: <0.01; ***: <0.001; ****: <0.0001.

### Transgene expression from lentiviral, adenoviral and plasmid vectors is inhibited by MORC3 and SETDB1

In addition to rAAV, other viruses such as lentivirus and adenovirus are used for clinical gene delivery applications (21). Interestingly, several recent studies suggested that MORC3 and SETDB1 can suppress the expression of retroviral elements (10, 19, 22). Consistent with previous studies, both MORC3 and SETDB1 KO increased lentivirus transduction rates and the MFI of encoded GFP across multiple lentiviral dilutions (Figure 5A and 5B). This effect was overall comparable as for rAAV, however, in contrast to rAAV, deletion of MORC3 had a stronger effect on lentiviral transgene expression (e.g. ~12x increase in MFI at 1:3 lentivirus dilution) than deletion of SETDB1 (~8x increase in MFI). RT-qPCR of lentiviral mRNAs showed that transgene expression changes were higher than for rAAV2 in MORC3 KO and SETDB1 KO cells suggesting stronger cellular restriction (Figure S4B). During infection with GFP-expressing adenovirus 5 (AdV5-GFP), the percentage of GFP+ cells and the MFI of GFP were also increased in MORC3 and SETDB1 KO relative to WT cells (Figure 5C and 5D). Moreover, transfection of a GFP-expressing plasmid also led to an increased MFI in MORC3 and SETDB1 KO cells relative to A549 WT cells, although the overall percentage of GFP+ cells did not change (Figure 5E and 5F). By contrast, expression of a fluorescent protein from transfected mRNA-liposomes was unchanged in MORC3 and SETDB1 KO relative to WT cells (Figure 5G and 5H). Together, these data suggest MORC3 and SETDB1 broadly restrict foreign DNA elements, such as viral vectors and plasmid DNA.

**Figure 5.**
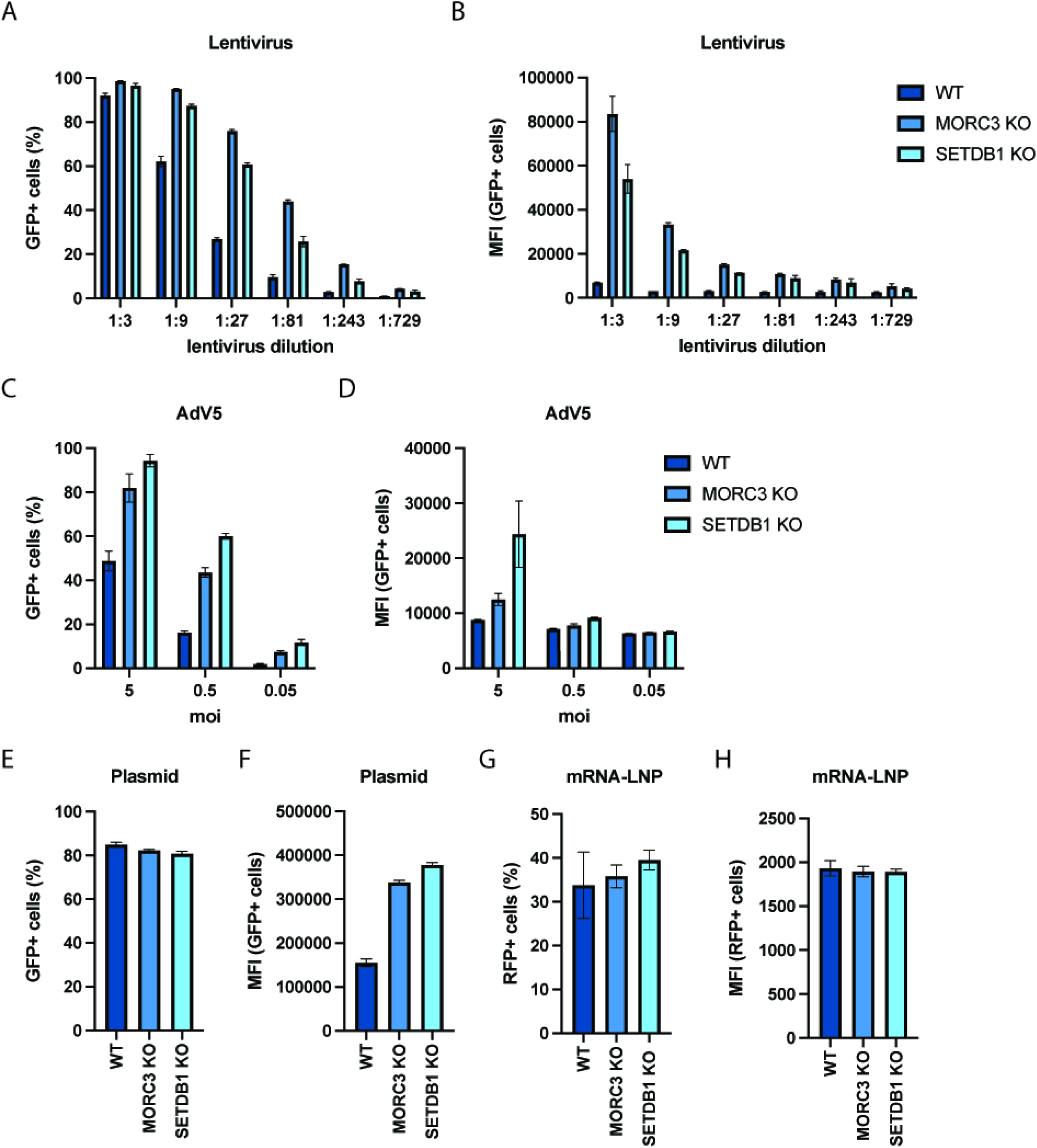
MORC3 and SETDB1 repress transgene expression from lentiviral, adenoviral and plasmid vectors. (A) Percentage or (B) MFI of WT, MORC3 KO or SETDB1 KO A549 cells transduced with different dilutions of a GFP-expressing lentivirus. Fluorescent cells were measured by flow cytometry 3 days post-transduction. (C) Percentage or (D) MFI of WT, MORC3 KO or SETDB1 KO A549 cells infected with different moi of a GFP-expressing adenovirus 5 (AdV5-GFP) vector. Cells were analyzed by flow cytometry at 24 hpi. (E) Percentage or (F) MFI of WT, MORC3 KO or SETDB1 KO A549 cells transfected with a GFP encoding plasmid and analyzed by flow cytometry 24h post-transfection. (G) Percentage or (H) MFI of WT, MORC3 KO or SETDB1 KO A549 cells transfected with in-vitro transcribed RFP mRNA and analyzed by flow cytometry 24h post-transfection. All data are shown as the mean ± s.d. from biological triplicates.

### Knockdown of MORC3 and SETDB1 in primary human fibroblasts leads to increased transgene expression

Finally, we explored whether inhibition of MORC3 or SETDB1 activity also affected rAAV transgene expression in untransformed, primary human cells. For this purpose, we performed siRNA knockdown in BJ fibroblasts. Transfection with MORC3 or SETDB1 siRNAs prior to rAAV2-tdTomato transduction increased the percentage of tdTomato+ cells relative to cells transfected with a non-targeting siRNA from 10% to 54% for MORC3 knockdown and to 18% for SETDB1 knockdown (Figure 6A). RT-qPCR confirmed successful knockdown of MORC3 and SETDB1 mRNA levels, although residual expression may still restrict rAAV transgene expression to a certain degree (Figure 6B). Overall, this suggests that the rAAV restriction factors identified in the genome-wide CRISPR screens could have a physiologically relevant function in inhibiting rAAV-based therapies in clinical settings. Additional validation of screen hits beyond FANCA, MORC3 and SETDB1 is needed as well as confirmation in additional primary cells and in vivo models.

**Figure 6.**
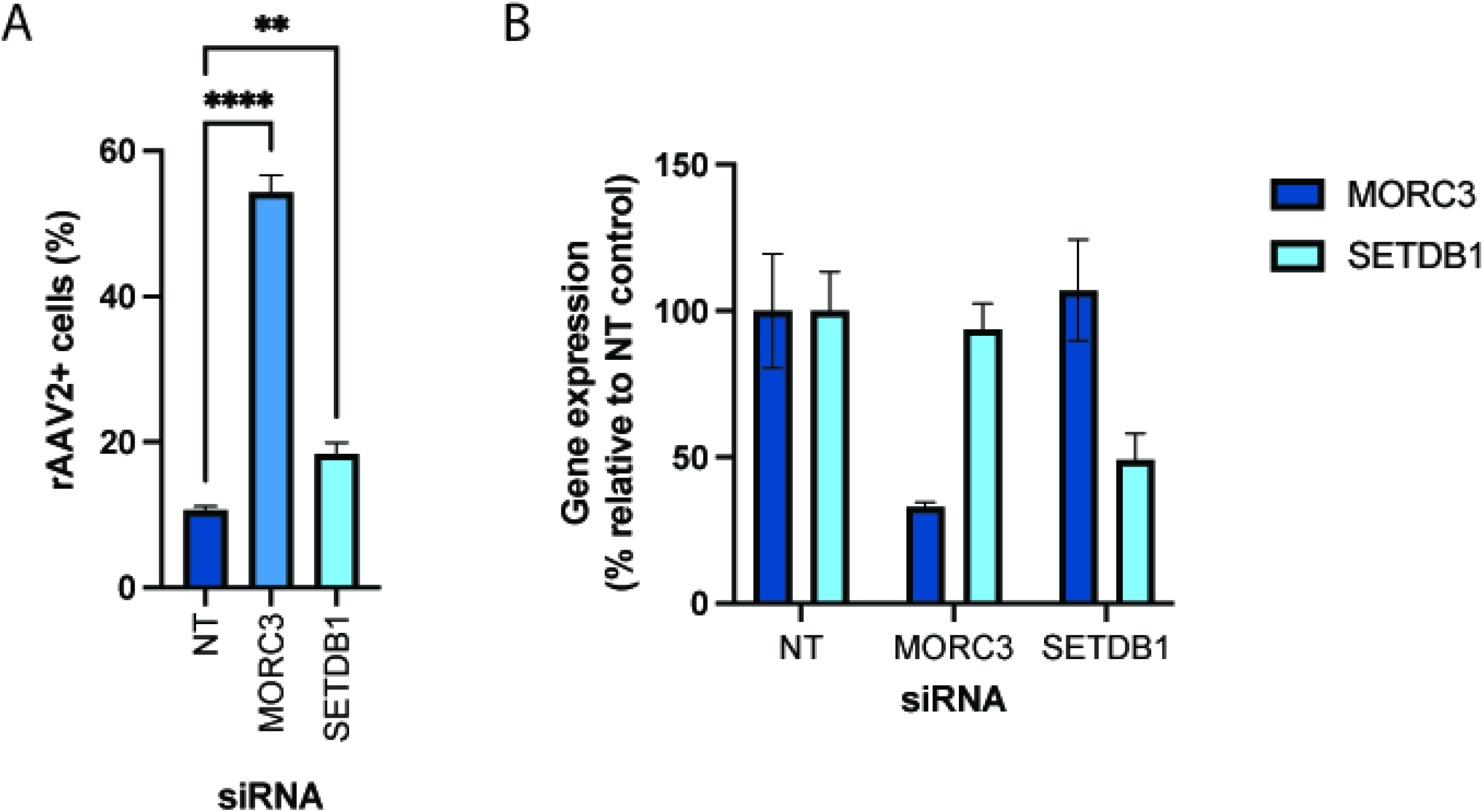
Knockdown of MORC3 and SETDB1 in primary human fibroblasts increases rAAV transgene expression. (A) Percentage of rAAV2-tdTomato positive BJ fibroblasts that were transfected with a non-target siRNA (NT) or siRNAs against MORC3 and SETDB1 4 days prior to rAAV transduction. Cells were analyzed by flow cytometry 36h post-transduction. Data are shown as the mean ± s.d. from biological triplicates. P-values were determined by ANOVA with post-hoc Tukey’s test and are defined as follows: ns: non-significant; *: <0.05; **: <0.01; ***: <0.001; ****: <0.0001. (B) Quantification of MORC3 and SETDB1 mRNA levels 4 days after siRNA transfection. Data are shown as the mean ± s.e.m. from biological triplicates.

## DISCUSSION

Our study provides a comprehensive picture of the cellular factors that limit gene expression from AAV vectors in human cells. The majority of the identified host genes have known functions in DNA damage sensing and repair, chromatin remodeling and transcriptional regulation. The most prominent hit across both CRISPR screens was the FA pathway, which is critical for the resolution of interstrand crosslinks or stalled replication forks (7). It is plausible that the ITRs in the AAV genome mimic interstrand crosslinks or stalled replication forks and are thus recognized by the FA machinery, which subsequently may impede second-strand synthesis of or transcription from the rAAV genome. Interestingly, another study found that deletion of FANCM led to enhanced AAV template-mediated homologous recombination (23). This further supports that the FA machinery engages with AAV genomes to interfere with both transgene expression as well as AAV template-mediated DNA repair. Other DNA damage response genes enriched in our dataset, e.g. the ATM kinase and the MRN complex (consisting of MRE11A/RAD50/NBN), have been shown to interact with the ITRs in the AAV genome and suppress transgene expression (24–26). Interestingly, studies with other DNA viruses suggested pro-viral effects for the FA pathway, which could be due to distinct differences in their genome structure and life cycle (27–30). Future studies are warranted to mechanistically investigate how the FA pathway interferes with the rAAV vectors and whether FA components directly bind to the AAV genome.

Moreover, we found that knockout of the HUSH-associated histone methyltransferase SETDB1 substantially increased transgene expression from rAAV and other viral vectors, such as lentivirus and adenovirus. However, this effect may be partially or largely due to an increase of intracellular rAAV genome copies. The exact mechanism how SETDB1 regulates rAAV genome stability and whether it additionally regulates transcriptional activity from vector genomes need to be studied further. In mammalian cells, SETDB1 is recruited by the HUSH complex to regulate heterochromatin and has been described to mediate silencing of human immunodeficiency virus (HIV) and other retroviral sequences by depositing repressive H3K9me3 histone marks (10, 11, 14, 31, 32). Importantly, SETDB1 is able to silence both integrated and unintegrated retroviral DNA (10, 14). rAAV genomes typically exist as unintegrated episomes and are thought to be bound by histones (33). Recently, it was also shown that the HUSH machinery recognizes and silences a broad range of long, intronless transgenes (34). SETDB1 and the HUSH complex may thus be major contributors to rAAV chromatinization and silencing, and potentially tagging heterochromatinized episomes for degradation. The crucial role for SETDB1 and the HUSH complex is further underscored by the enrichment of other factors in our screens. ATF7IP interacts with SETDB1 and protects it from proteasomal degradation (13). The DNA-binding protein ZNF638 (also known as NP220) is responsible for recruitment of the HUSH complex to unintegrated murine retroviral DNA and may also act similarly on rAAV genomes (14). MORC2 was shown to to mediate transgene silencing and maintain chromatin compactness upon recruitment by the HUSH complex (12). Finally, the role of the HUSH complex as a rAAV restriction factor was also corroborated by a recent study. Consistent with our data, it showed that knockout of HUSH components resulted in increased rAAV transgene levels and that deletion of ZNF638 (NP220) decreased repressive H3K9me3 histone marks on AAV genomes (35).

In addition to the HUSH-associated MORC2, MORC3, another Microchidia CW-type zinc finger protein, was the top hit in the A549 screen and strongly enriched in the K562 screen. MORC proteins are highly conserved, reside in the nucleus, contain a GHKL ATPase domain, and have been linked to heterochromatin condensation and gene silencing (17, 36). Previously, MORC3 was shown to restrict herpesviruses and endogenous retroviruses (ERVs) (18, 19). Mechanistic studies revealed that MORC3 bound to ERV sequences and that knockout of MORC3 resulted in ERV de-repression, reduced H3K9me3 levels and increased chromatin accessibility (19). According to our results, we propose that MORC3 is a critical rAAV restriction factor that silences rAAV genomes through possible chromatin regulation.

Additional screen hits have also been linked to chromatin regulation and gene silencing. The ATRX/DAXX complex is responsible for deposition of H3.3 histones at heterochromatic regions and has been shown to associate with both SETDB1 and MORC3 (19, 37, 38). Moreover, the CHAF1 complex, together with the histone chaperon ASF1, delivers newly synthesized H3/H4 dimers to replication forks and was recently shown to mediate silencing of unintegrated HIV-1 DNA (39, 40). Overall, these chromatin modifiers with described roles in foreign DNA silencing and may therefore also be involved in rAAV repression.

Lastly, the SMC5-SMC6 complex, which is critical for genome stability and DNA repair (41), has emerged as a component of antiviral responses. It has recently been linked to transcriptional restriction of numerous DNA viruses, such as hepatitis B virus, unintegrated retroviral DNA, papillomavirus and Epstein-Barr virus (15, 16, 42, 43). Our data suggests that SMC5-SMC6 can also potently inhibit rAAV vectors. During DNA repair, this complex is recruited to DNA lesions by FAM178A (also known as SLF2) and SLF1 (44). A recent preprint demonstrated that SIMC1, a top hit in both CRISPR screens and largely with unknown function, forms a distinct complex with FAM178A to localize SMC5/SMC6 to viral replication centers (45). Therefore, it is plausible that rAAV is repressed by recruitment of the SMC5-SMC6 complex via FAM178A/SIMC1.

Overall, the identification of multiple components of specific cellular pathways in our CRISPR screens indicates that these are high-confidence candidates for rAAV restriction factors. Of note, two previous siRNA screens for regulators of rAAV transduction in HeLa cells showed enrichment of a subset of the same restriction factors as in our CRISPR screens, including ATF7IP, CHAF1A, DAXX, NPAT (a positive regulator of ATM) and MORC3 (previously named ZCWCC3) in one study, as well as ATF71P, CHAF1A, SETDB1, NPAT and the transcription factor MTF1 in another study (46, 47). Thus, our work corroborates some previously identified candidate restriction factors and expands the network of cellular factors that interfere with rAAV transgene expression.

## MATERIAL AND METHODS

### Cell Culture

A549 and K562 were obtained from ATCC and cultured in DMEM (Gibco) supplemented with 10% fetal bovine serum (Omega Scientific), penicillin/streptomycin (Gibco), non-essential amino acids (Gibco) and L-glutamine (Gibco) at 37C and 5% CO2. HEK293ft cells were obtained from Thermo Scientific and cultured as described above. WT, FANCA KO and trans-complemented FANCA KO UM-SCC1 cells, WT and FANCA KO mouse ear fibroblasts, and FANCA-deficient patient-derived 974 cells with and without FANCA complementation were kindly provided by the Fanconi Anemia Research Fund at Oregon Health & Science University and cultured as described above. Normal human BJ fibroblasts were obtained from ATCC (CRL-2522) and cultured in Knockout DMEM (Gibco) supplemented with 15% fetal bovine serum, 16.5% Medium 199 (Gibco), penicillin/streptomycin, and L-glutamine at 37C and 5% CO2.

### Viruses

The following rAAV vectors were obtained from the UNC School of Medicine Gene Therapy Center Vector Core: AAV serotype 2 expressing GFP under a chimeric CMV–chicken ß–actin (CBA) promoter (rAAV2-GFP), AAV serotype 2 expressing tdTomato under a CMV immediate enhancer/β-actin (CAG) promoter (rAAV2-tdTomato), AAV serotype 1, 5 or 6 expressing GFP under a CMV promoter (rAAV1/5/6-GFP), and self-complementary AAV serotype 2 expressing GFP under a CMV promoter (scAAV2-GFP). Lentivirus expressing GFP was produced by co-transfection of HEK293ft cells with the following plasmids: plenti-CMV-Puro-DEST (Addgene #17452, gift from Eric Campeau & Paul Kaufman) where GFP cDNA was cloned into the EcoRV site using NEBuilder HiFi DNA Assembly Master Mix; pCMV-dR8.2 dvpr (Addgene, #8455, gift from Bob Weinberg); pCMV-VSV-G (Addgene, #8454, gift from Bob Weinberg); and pAdVAntage (Promega). Supernatants were collected 48h post-transfection, filtered and stored at −80C until use. Adenovirus Serotype 5, Clone Ad5-CMV-hACE2/RSV-eGFP (NR-52390) was kindly provided through BEI Resources.

### CRISPR screens

A549 cells were stably transduced with lentivirus from lentiCas9-Blast (Addgene #52962, gift from Feng Zhang) and subsequently selected using blasticidin. Next, a total of 240 million A549-Cas9 cells were transduced with the lentivirus of the human GeCKO v2 library (Addgene #1000000049, gift from Feng Zhang) (48) at a moi of 0.4 and selected using puromycin for 7 days. 60 million cells of the A549 CRISPR KO library were collected for genomic DNA (gDNA) extraction to assess the reference gRNA distribution. Another 60 million cells were transduced with rAAV2-GFP at a moi of 10,000 vg/cell, which resulted in a ~45-50% transduction rate. The top 10-15% GFP+ cells were sorted 3 dpi and replated. After 7 days, the A549 cells were re-transduced with 10,000 vg/cell and the top 10-15% GFP+ cells were sorted again. A total of ~8 million cells was collected for gDNA extraction.

K562-dCas9-KRAB cells were a gift from Jonathan Weissman. 240 million cells were transduced with lentivirus from the Human Genome-wide CRISPRi-v2 (Addgene #83969, also gift from Jonathan Weissman) (49) at a moi of 0.4 using spin-inoculation (1000g, 33C, 2h) in 6-well plates. Subsequently, cells were selected using Puromycin for 5 days. 100 million K562 CRISPRi library cells were collected for gDNA extraction to assess the initial gRNA distribution. Another 60 million cells were spin-infected (1000g, 33C, 2h) with AAV2-GFP at a moi of 50,000 vg/cell. The top 15-20% GFP+ cells (out of a total %GFP+ of 60%) were sorted and then continued to be cultured. After 7 days, another round of spin-infection with AAV2-tdTomato and FACS of the top 10-15% tdTomato+ cells were performed. A total of 16 million cells were collected for gDNA extraction.

For both A549 and K562 cells, gDNA was isolated using the Qiagen DNA Blood Maxi kit (for reference cells) or multiple QIAamp DNA Mini kit columns (for sorted cells). CRISPR gRNA encoding DNA sequences were amplified in a two-step nested PCR using KAPA HiFi HotStart ReadyMixPCR Kit (Kapa Biosystems). In the first PCR step, 36-48 reactions and 12 reactions containing 6 μg gDNA were set up for the reference and sorted samples, respectively, and amplified for 16 cycles. Reactions for each sample were pooled and mixed. In the second PCR step, 4 reactions containing 5 μl PCR1 product were amplified for 12 cycles using indexed primers. PCR products were gel purified using QIAquick Gel Extraction Kit and sequenced on an Illumina NextSeq 500 using custom sequencing primers. Primers sequences are listed in Table S4.

For analysis, demultiplexed FASTQ files were aligned to gRNA reference table and enrichment of each gRNA was calculated by comparing the relative abundance in the selected and unselected cell population. Gene enrichment analysis was performed using MaGeCK (50). For STRING analysis (51), the top 100 enriched genes from each CRISPR screen were used as input and gene interactions were computed using a STRING confidence score of 0.4. MaGeCK scores were overlaid using Cytoscape 3.9.0 (52) and only the major gene network is displayed in Figure 1.

### Generation of KO cell lines

DNA oligos (Integrated DNA Technologies) containing gRNA sequences complementary to MORC3 or SETDB1 genomic loci were annealed and ligated into pX458 (Addgene #48138, gift from Feng Zhang). A549 cells were transfected with pX458 constructs using Lipofectamine 3000 and two days later GFP-positive cells were single-cell sorted into 96-well plates containing complete media. After several weeks, gDNA was isolated from obtained cell clones using QuickExtract (Lucigen), the gRNA-targeted sites were PCR-amplified, PCR products Sanger-sequenced and sequences aligned to reference sequences using Geneious Prime. A list of all used gRNA oligo and genotyping primer sequences can be found in Table S4.

### Lentiviral complementation of KO cells

For generation of trans-complemented cell lines, lentivirus was generated as described above using plenti-CMV-Puro-DEST containing MORC3 (OriGene Technologies, RC210530) or SETDB1 (OriGene Technologies, RC226620) cDNA sequences. KO cells were transduced in presence of polybrene and subsequently selected using Puromycin.

### Flow cytometry

Cells were plated in 96-wells and infected the next day. At indicated timepoints, cells were trypsinized and analyzed by flow cytometry using a Cytoflex S flow cytometer (Beckman Coulter). At least 4,000 cells were recorded per sample and cells were gated based on FSC/SSC, FSC-H/FSC-A (singlets) and finally PE (tdTomato) or FITC (GFP) to determine the percentage of fluorescent (infected) cells using FlowJo 10. Moreover, the median fluorescence intensity (MFI) of the fluorescent cells was determined using FlowJo 10. In all experiments, uninfected cells served as negative control for gate setting.

### Quantification of intracellular AAV genome copies

UM-SCC1 or A549 cell lines were plated in 12-well plates and transduced with rAAV2-tdTomato at a moi of 10,000 vg/cell the next day. 48h post-transduction cells were washed with PBS 3 times, trypsinized and pelleted by centrifugation. Cellular and rAAV DNA was extracted from cell pellet using the DNeasy Blood and Tissue kit (Qiagen). 2 μl of purified DNA was used as input to quantify AAV genome copies via qPCR on a Bio-Rad CFX96 Touch system. AAV genomes copies were determined by amplification with primers targeting the AAV2 ITR region and normalized to the cellular RPPH1 locus copy number using IDT PrimeTime qPCR primers. QPCR primer sequences can be found in Table S4.

### RT-qPCR

For quantification of transgene expression, AAV- or lentivirus transduced cells were harvested using the Power SYBR Green Cells-to-CT kit (Invitrogen). After reverse transcription, quantitative PCR was performed on a Bio-Rad CFX96 Touch system and viral transgene levels (GFP or tdTomato) were normalized to cellular 18S levels. For quantification of siRNA knockdown of MORC3 and SETDB1 mRNA levels, cells were harvested 4 days post siRNA transfection using the Power SYBR Green Cells-to-CT kit (Invitrogen). IDT PrimeTime qPCR Primers for MORC3 and SETDB1 were used and expression levels were normalized to 18S. QPCR primer sequences can be found in Table S4.

### Cell proliferation assay

WT, MORC3 KO and SETDB1 KO cells were seeded in 96-wells at a density of 8,000 cells per well. At 24, 48 and 72h post-plating Cell Titer Glo 2.0 solution (Promega) was added on top of the cell media and incubated for 10min. After that, 100 μl were transferred to a white 96-well microplate and luminescence was measured using on an EnVision plate reader (PerkinElmer).

### siRNA transfection

ON-TARGETplus Non-targeting Control Pool, ON-TARGETplus Human MORC3 siRNA SMART POOL and ON-TARGETplus Human SETDB1 siRNA SMART POOL were obtained from Horizon Discovery.

10,000 BJ fibroblasts were plated in 96-wells to reach ~50% confluency by the next day. Then, cells were transfected with siRNA pools using 0.3 μl RNAiMAX reagent (Invitrogen), 30 nM siRNA and 20 μl OptiMEM (Gibco) per 96-well. 4 days post-transfection, cells were transduced with rAAV and rAAV transgene expression was quantified by flow cytometry 36h post-transduction. In parallel, transfected but untransduced cells were harvested for analysis of mRNA knockdown.

## Supporting information

Combined Supplemental Figures

## DATA AVAILABILITY

The raw sequencing data for the A549 CRISPR KO and K562 CRISPRi screens are available on NCBI under the BioProject ID PRJNA852611.

## ACKNOWLEDGMENTS

We thank Drs. Nicole Paulk (UCSF), Jan Carette (Stanford University) and James Zengel (Stanford University) for helpful discussions; the Biohub Genomics Platform for help with sequencing of the CRISPR screen samples; the UNC Gene Therapy Center for providing rAAV vectors; Drs. Jonathan Weissman (UCSF) and Feng Zhang (Broad Institute) for providing the CRISPRi and CRISPR KO library plasmids; and Dr. Leslie Wakefield and the Fanconi Anemia Research Fund at Oregon Health & Science University for providing FANCA KO and trans-complemented cell lines.

## AUTHOR CONTRIBUTIONS

AMN and ASP performed all experiments and analyzed data. ASP designed the study and wrote the manuscript.

## DECLARATION OF INTERESTS

The authors have declared no competing interest.

## Notes

### Summary of Updates

This revision contains new data (Fig. 2G, 3I, 5E, 5F, 6) as well as additional discussions/clarifications in the text.

