## Supplementary material for "Genome-scale analysis of cellular restriction factors that inhibit transgene expression from adeno-associated virus vectors": Combined Supplemental Figures

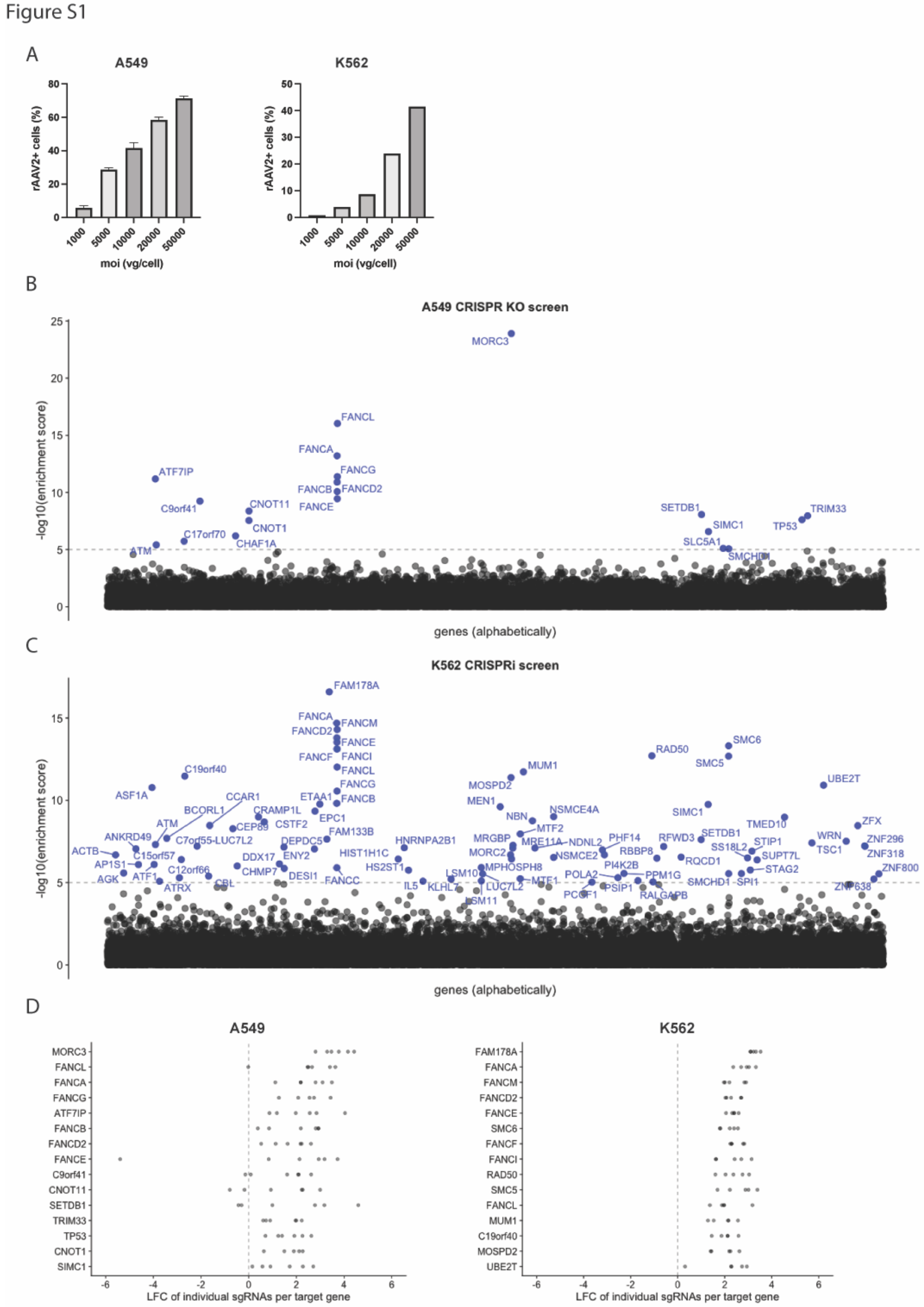


**Figure S1. rAAV CRISPR screen optimization and analysis.**

(A) Moi titration of rAAV2-GFP on A549 and K562 cells. Infection rate was measured by flow cytometry 24hpi.
(B) MaGeCK analysis of the A549 CRISPR KO screen. Genes with -log10 (enrichment score) > 5 are highlighted.
(C) MaGeCK analysis of the K562 CRISPRi screen. Genes with -log10 (enrichment score) > 5 are highlighted.
(D) Log-fold changes (LFC) of individual gRNAs of the top 15 scoring genes from the A549 and K562 CRISPR screens.

Figure S2

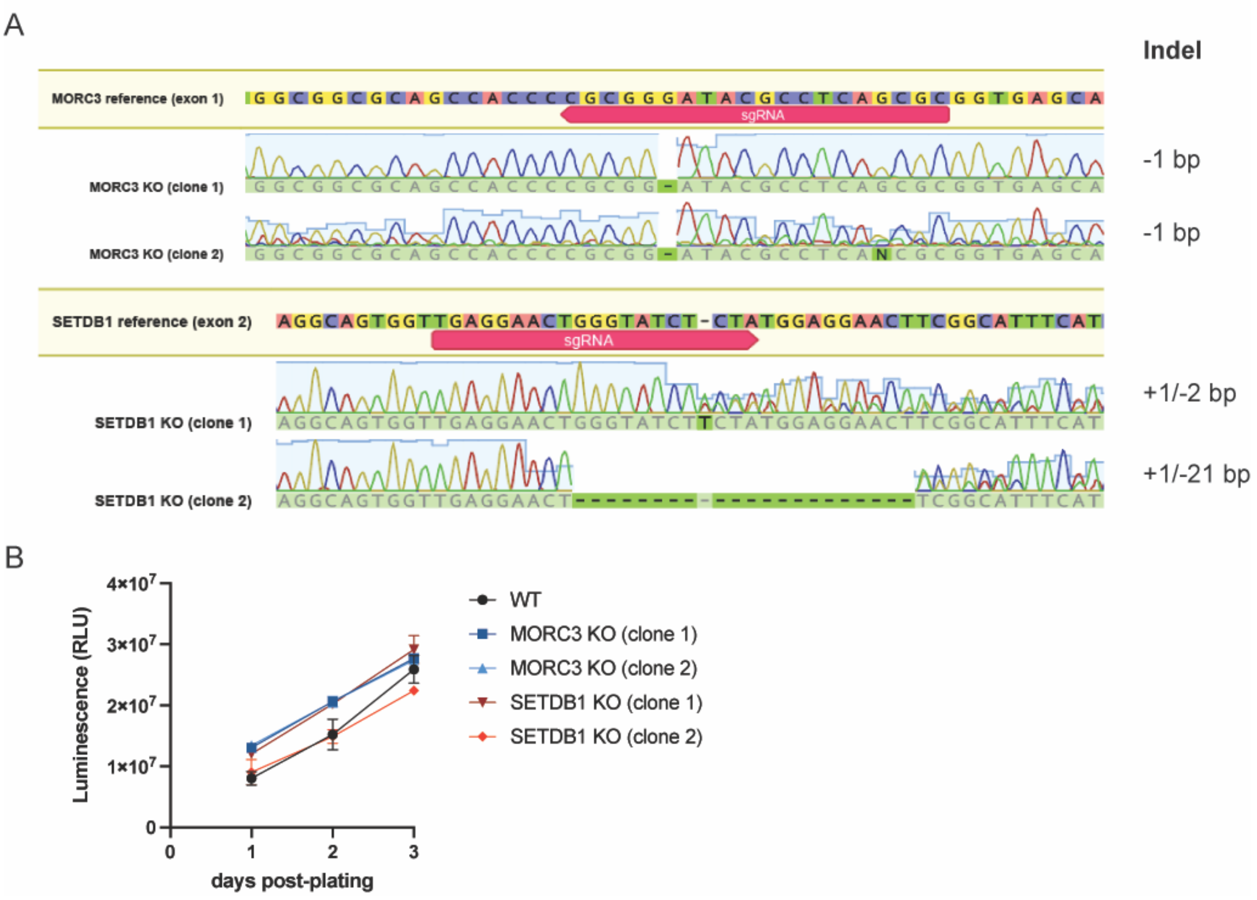


**Figure S2. Characterization of MORC3 and SETDB1 KO cell lines.**

(A) Genotyping data of clonal MORC3 and SETDB1 KO cell lines. gRNA-targeted genomic loci were amplified by PCR, products were Sanger-sequenced, and reads were aligned to reference sequences.

(B) Cell proliferation of WT and clonal KO cell lines. Cells were harvested at each timepoint and cell proliferation was measured by Cell Titer Glo assay. Data is displayed as mean ± s.d. from biological triplicates.

Figure S3

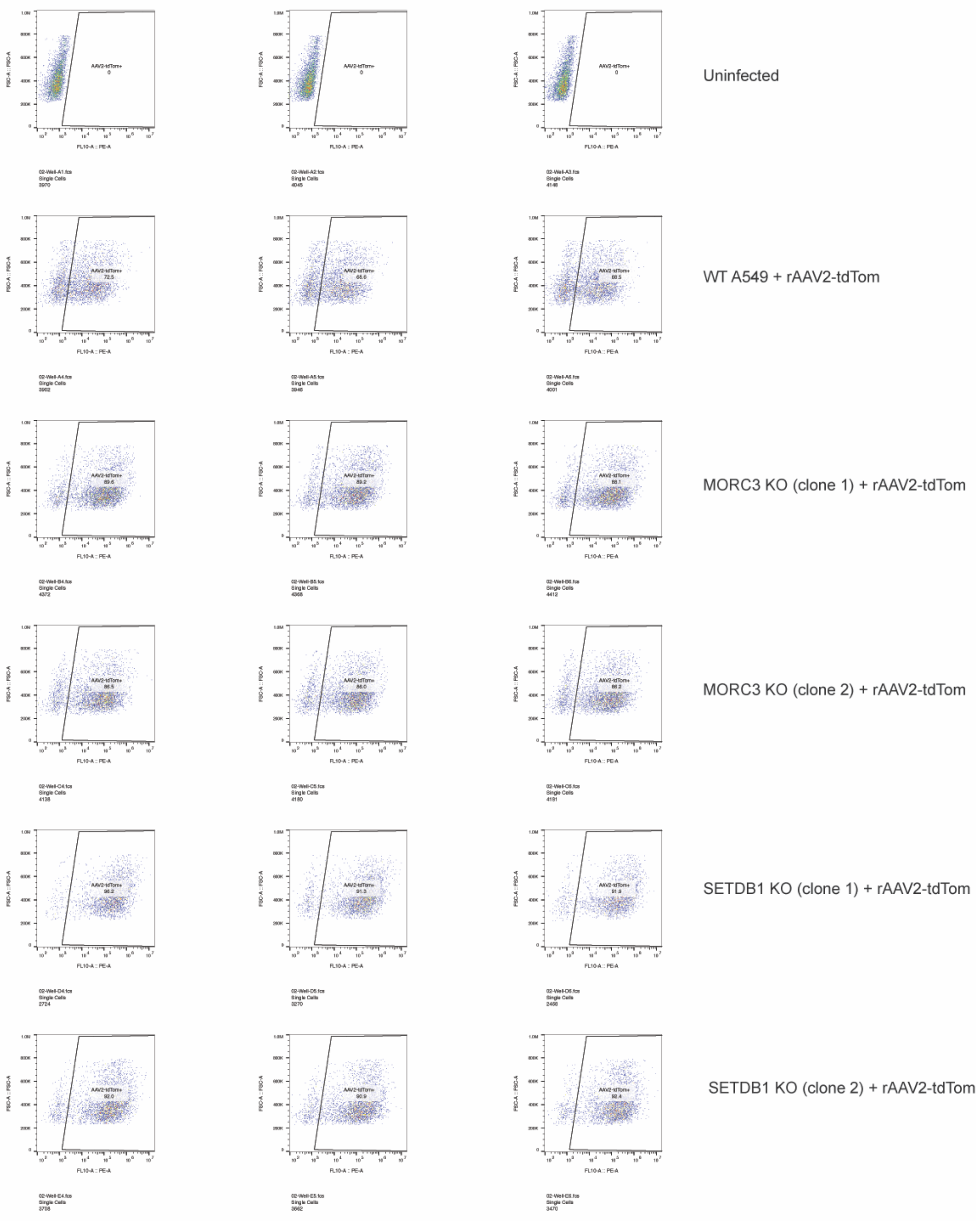


**Figure S3. Representative flow cytometry data for rAAV2-tdTomato infection of WT, MORC3 KO and SETDB1 KO A549 cells.** Cells were infected with tdTomato-expressing rAAV2 (moi=10,000) and measured by flow cytometry at 3dpi. Each panel represents one biological replicate. Summary data are displayed in Figure 3A and 3B.

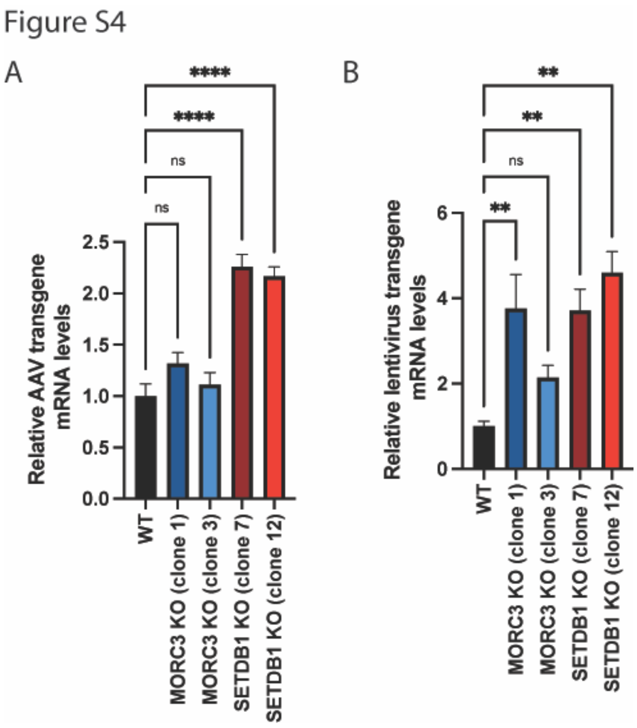


**Figure S4. Knockout of MORC3 and SETDB1 increases transgene mRNA levels from rAAV and lentiviral vectors.**

(A) rAAV2 tdTomato transgene mRNA levels in WT, MORC3 KO or SETDB1 KO A549 cells normalized to 18S ribosomal RNA levels.

(B) Lentiviral GFP transgene mRNA levels in WT, MORC3 KO or SETDB1 KO A549 cells normalized to 18S ribosomal RNA levels.

All data are shown as the mean ± s.e.m. from biological triplicates. P-values were determined by ANOVA with post-hoc Tukey's test and are defined as follows: ns: non-significant; *: <0.05; **: <0.01; ***: <0.001; ****: <0.0001.

**SUPPLEMENTAL TABLES**

**Table S1: MaGeCK analysis of A549 CRISPR KO screen for increased AAV2 transgene expression.**

**Table S2: MaGeCK analysis of K562 CRISPRi screen for increased AAV2 transgene expression.**

**Table S3: Functional enrichment analysis for biological processes, molecular function and cellular components of the STRING network containing the top 100 genes from the A549 and K562 CRISPR screens.**

**Table S4: DNA oligos used in this study.**
